# Host-derived lipoprotein is an environmental cue for asexual reproduction via a lipoprotein receptor homolog in a cestode

**DOI:** 10.64898/2026.07.17.739065

**Authors:** Taizo Saito, Kei Hayashi, Keisuke Nakagawa, Yasuhiro Takashima

## Abstract

Parasitic helminths undergo major changes in their host environment during their life cycles, including transitions between hosts, and respond by initiating specific developmental programs such as asexual reproduction. These programs are thought to be triggered by parasite recognition of host-derived environmental cues. However, the molecular identity of such host-derived cues and the mechanisms by which they are recognized remain unknown. Here, we investigated this mechanism in the cestode *Mesocestoides vogae*, which undergoes asexual reproduction in the intraperitoneal cavity of intermediate hosts such as mice. To mimic the intraperitoneal environment, serum was used as a proxy for conditions in peritoneal fluid in vitro. Mouse serum initiated asexual reproduction in vitro, whereas guinea pig serum did not. We found that the parasite establishes infection in the peritoneal cavity of guinea pigs but fails to proliferate. The marked difference in high-density lipoprotein (HDL) levels between mouse and guinea pig serum prompted us to test HDL, which initiated asexual reproduction. Knockdown of a putative HDL receptor homolog significantly reduced the frequency of asexual reproduction, supporting its role in the process. Together, these findings indicate that *M. vogae* uses HDL as a host-derived environmental cue through an HDL receptor homolog to trigger asexual reproduction. Here, we demonstrate that parasitic helminths can discriminate specific host-derived factors within complex host environments to initiate developmental programs.

**Significance Statement:** Parasitic helminths, including cestodes, undergo major developmental transitions in response to host environments, such as upon entry into specific organs. These transitions are thought to be triggered by host-derived factors, yet the identity of the host cues and the parasite molecules that sense them remain unknown. Here, we identify high-density lipoprotein as a host-derived cue that triggers asexual reproduction in *Mesocestoides vogae* and a candidate parasite receptor that senses this signal. Here, we provide molecular insight into how parasitic helminths sense environmental cues to trigger developmental switches.

## Introduction

Parasitic helminths, including cestodes and trematodes, undergo dramatic developmental changes, including marked alterations in behavior and morphology, as they transition between life-cycle stages within and between hosts (1–4). For example, trematode larvae undergo asexual reproduction and transformation within mollusks, their intermediate hosts, and subsequently emerge as motile cercariae that search for a second intermediate host or a definitive host outside the mollusks (1–4), illustrating how parasites adapt their behavior and morphology to specific host environments. These stage-specific transformations are critical for multiple aspects of the parasite life cycle, including propagation, stage conversion, and tissue colonization (1–4), yet the molecular mechanisms that trigger such developmental switches remain largely unknown.

The habitat of parasitic helminths is essentially confined to the host body (1–4), where environmental cues that regulate their developmental and behavioral transitions are encountered. These cues originate within the host, where larvae must detect host-derived factors to progress through the life cycle. Among these developmental events, asexual reproduction occurs primarily within the intermediate host. By increasing parasite numbers, this process is thought to confer adaptive advantages, including enhanced transmission success to their next host (2, 5) and improved competitive performance against other parasite species (6). Although many studies have examined the regulatory mechanisms underlying asexual reproduction in parasitic helminths (7–10), most have focused on regulation after reproduction has already been initiated. By contrast, the host-derived factors that trigger its onset have not yet been clearly identified in any parasitic helminth species.

*Mesocestoides vogae* is a cestode infecting carnivores as definitive hosts (11, 12), with larvae capable of asexual proliferation in intermediate hosts (11). In permissive intermediate hosts such as mice, larvae multiply extensively in the peritoneal cavity (11), whereas cultures in vitro allow long-term survival without initiating proliferation (13). This host-dependent pattern suggests that host-derived factors act as cues to trigger stage-specific larval expansion.

In the present study, we identified high-density lipoprotein (HDL) as a key host-derived cue. In addition, RNA interference (RNAi)-mediated knockdown of a gene encoding a putative HDL receptor in *M. vogae* resulted in impaired asexual reproduction. These findings provide molecular insight into how parasitic helminths sense a host-derived environmental cue and initiate developmental transitions.

## Results

### Mouse serum triggered the initiation of asexual reproduction in *Mesocestoides vogae* tetrathyridia

Tetrathyridia proliferate in the host peritoneal cavity by asexual reproduction under continuous exposure to peritoneal fluid (Fig. 1*A, B*). Asexual reproduction proceeds through scolex duplication, in which a single scolex gives rise to two daughter scolices (Fig. 1*C*). Expansion of the separating region between the two scolices then leads to separation into two independent tetrathyridia. Accordingly, scolex duplication was defined as the initiation of asexual reproduction.

**Figure 1.**
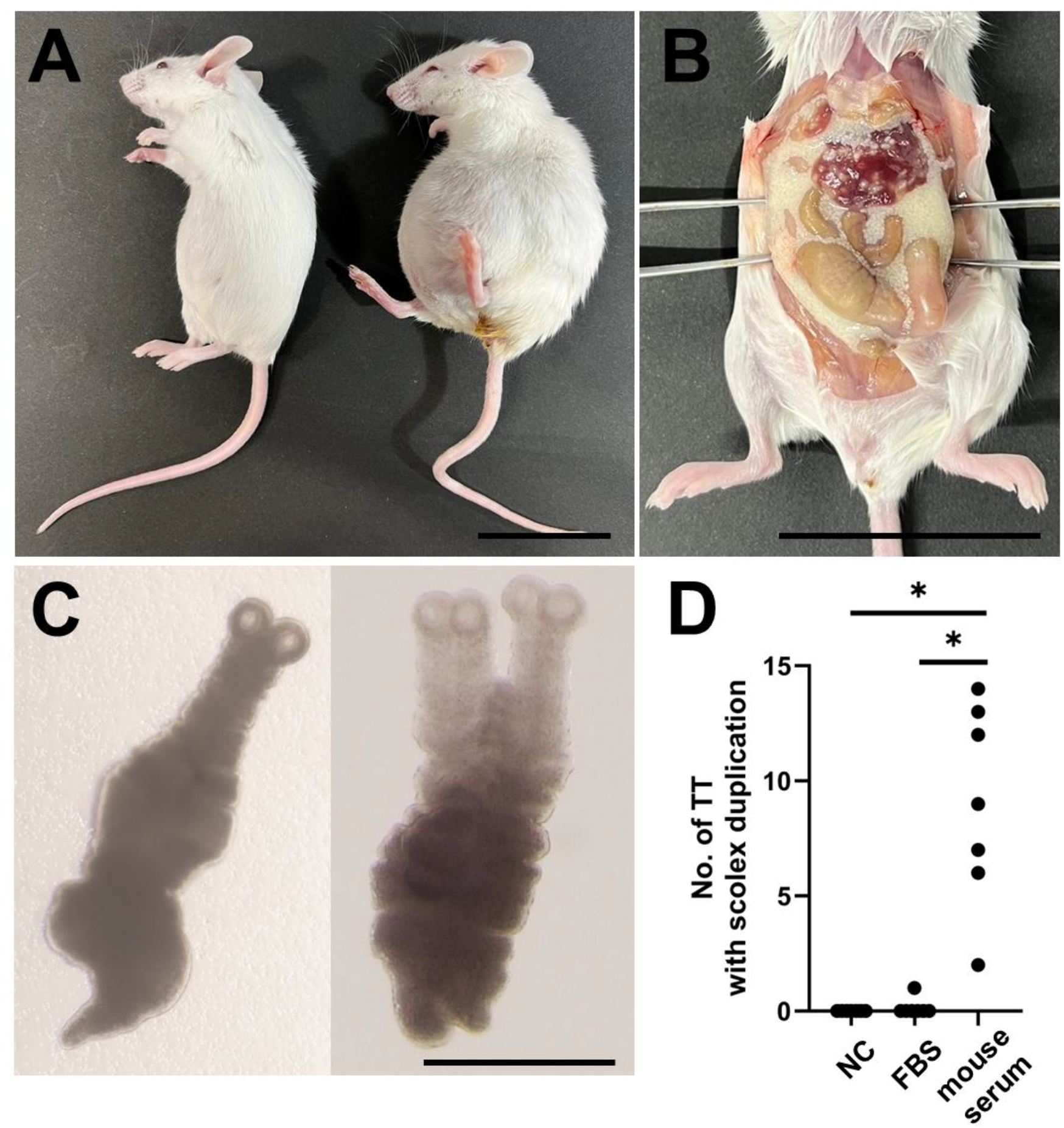
Mouse serum induces scolex duplication in *Mesocestoides vogae* tetrathyridia. (A) Uninfected mouse (left) and infected mouse (right). (B)Tetrathyridia residing in the peritoneal cavity of mice. The white granular material filling the interorgan spaces consists entirely of tetrathyridia. (C) Tetrathyridium with a single scolex (left) and a tetrathyridium in which duplication has occurred, resulting in two scolices (right). (D) Effect of serum supplementation on the initiation of scolex duplication in vitro. Single-scolex tetrathyridia were cultured in basic medium supplemented with fetal bovine serum (FBS), mouse serum, or without additional serum supplementation (NC). Each dot represents an independent experimental replicate. Significance was evaluated using a two-sided Wilcoxon signed-rank test with Holm’s correction for multiple comparisons. \**P* < 0.05. Scale bars = 5 cm (A, B) and 500 μm (C).

We hypothesized that peritoneal fluid contains factors that initiate this process. Because peritoneal fluid is largely derived from blood, the factors of interest are likely blood-borne (14). For this reason, host serum was used to examine these factors. The subsequent analyses therefore focus on serum-derived components.

To test whether serum contains factors that initiate asexual reproduction, tetrathyridia were cultured in medium supplemented with either mouse or fetal bovine serum; mice are a known permissive host (11), whereas cattle are presumed to be non-permissive. Under control conditions without additional serum supplementation, or in the presence of fetal bovine serum, scolex duplication was rarely observed (Fig. 1*D*). By contrast, supplementation with mouse serum consistently induced scolex duplication in all experimental replicates (Fig. 1*D*).

### Narrowing down the factor triggering the initiation of asexual reproduction

To narrow down the serum factor responsible for initiating scolex duplication, mouse serum was fractionated by ultrafiltration into two fractions based on molecular size, and each fraction was tested separately in tetrathyridial culture. The retentate fraction (upper; approximately 60 kDa) induced scolex duplication in significantly more worms than the filtrate fraction (lower) (Fig. 2*A*), indicating that the activity resides in components larger than approximately 60 kDa. We next exploited differences in host permissiveness to narrow the candidate factor further.

**Figure 2.**
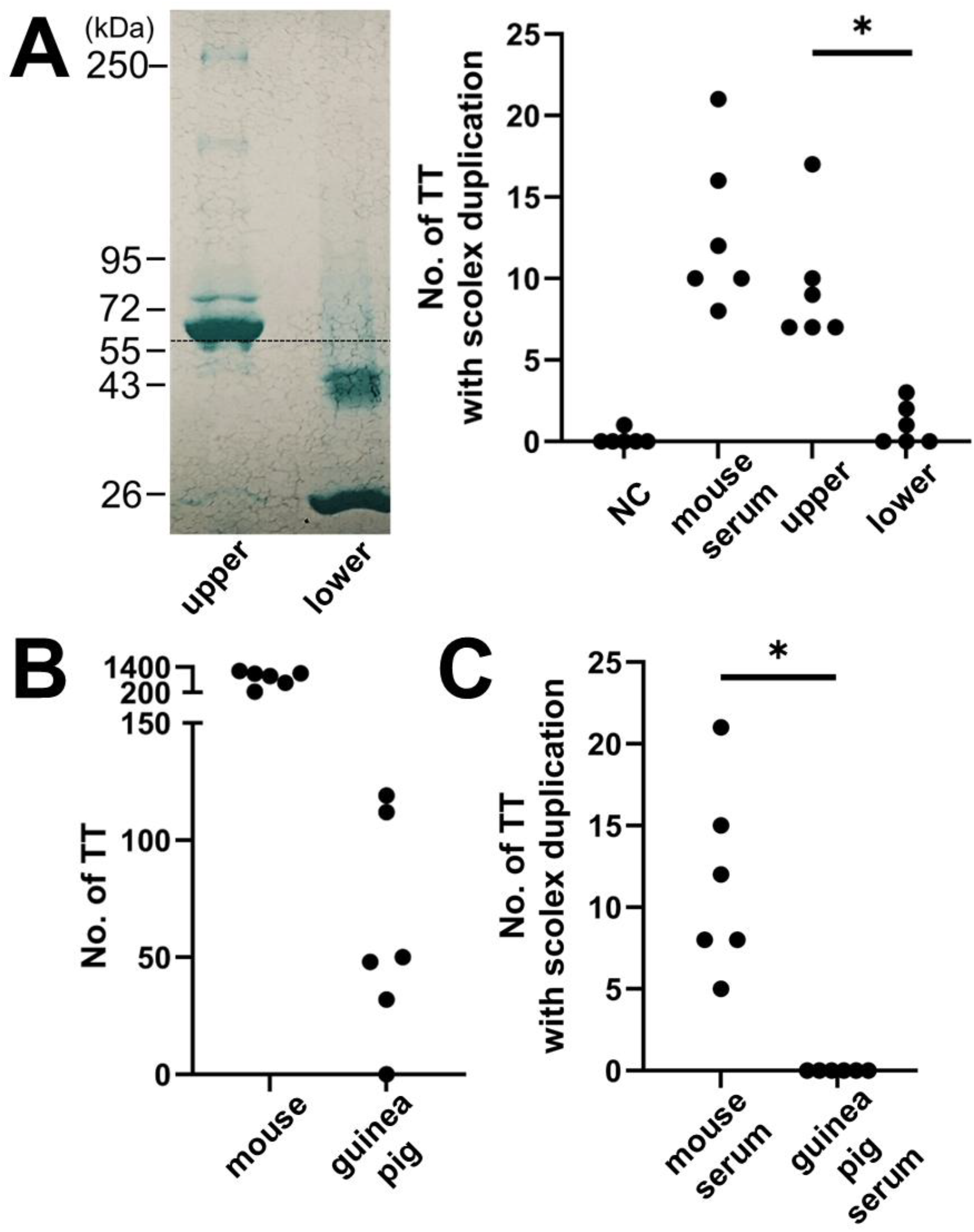
Characteristics of host serum factors initiating scolex duplication. (A) Mouse serum was fractionated by ultrafiltration into retained (upper) and flow-through (lower) fractions, which were tested separately in tetrathyridial culture. (B) Tetrathyridia were inoculated into the peritoneal cavities of mice and guinea pigs, and parasite numbers were determined 1 month after infection. (C) Tetrathyridia cultured with guinea pig serum after 7 days in vitro. Significance was evaluated using a two-sided Wilcoxon signed-rank test. \**P* < 0.05.

Tetrathyridia were inoculated into the peritoneal cavities of guinea pigs and mice, and the number of recovered parasites was determined 1 month later. Tetrathyridia recovered from guinea pigs showed no obvious differences in motility or morphology compared with those recovered from mice (Movies S*1* and S*2*, respectively), whereas parasite numbers were lower than the initial inoculum in all animals (Fig. 2*B*), indicating that guinea pigs represent a non-permissive host in which tetrathyridia survive but do not proliferate. Consistent with these findings, guinea pig serum supported parasite survival in vitro but failed to induce scolex duplication after 7 days of culture (Fig. 2*C*).

### Host-derived HDL triggers the initiation of asexual reproduction

Based on these results, we considered candidate factors larger than approximately 60 kDa whose concentrations differ among mice and guinea pigs. HDL satisfied these criteria. Guinea pigs have low plasma HDL concentrations (15). To test whether HDL could trigger scolex duplication, HDL isolated from mouse serum was added to the culture medium. Scolex duplication was observed in all experimental replicates (Fig. 3*A*). To assess the abundance of HDL in peritoneal fluid, fluid was collected from a subset of infected mice in which sufficient fluid had accumulated for collection. As a surrogate marker of HDL abundance, we measured HDL-cholesterol (HDL-C), a component of HDL that has been reported to correlate with HDL particle levels (16). HDL-C was detected in all peritoneal fluid samples at approximately half the corresponding serum concentration (Table S*4*).

**Figure 3.**
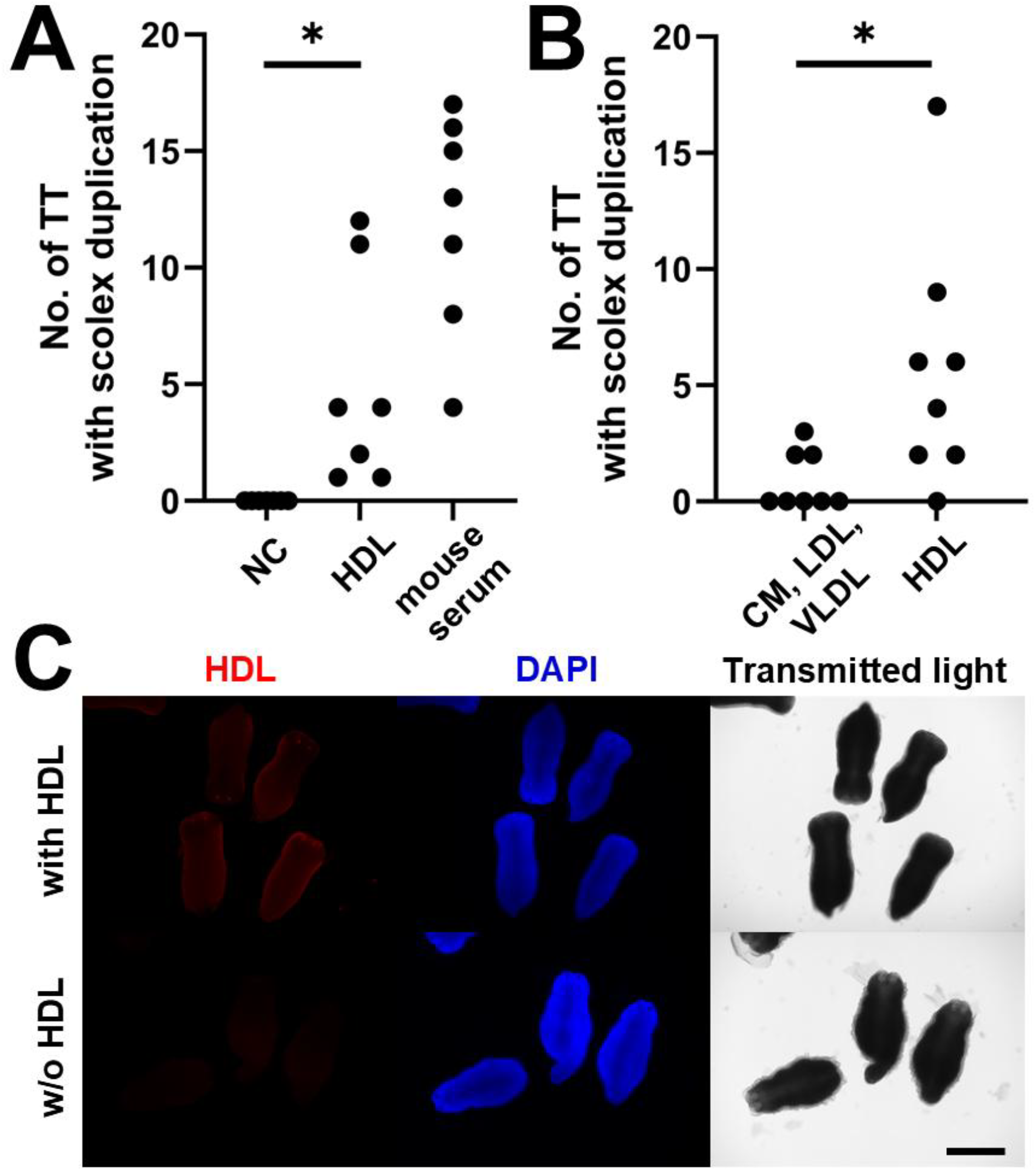
High-density lipoprotein (HDL) acts as a host-derived environmental cue inducing scolex duplication. (A) HDL isolated from mouse serum was added to tetrathyridial culture medium. (B) Serum lipoproteins were separated by ultracentrifugation into HDL and non-HDL fractions and tested in tetrathyridial culture. (C) Tetrathyridia cultured with fluorescently labeled HDL. Significance was evaluated using a two-sided Wilcoxon signed-rank test. \**P* < 0.05. Scale bar = 500 μm.

To determine whether lipoproteins other than HDL, including low-density lipoprotein (LDL), very low-density lipoprotein (VLDL), and chylomicrons (CM), could also induce this process, serum lipoproteins were separated by ultracentrifugation into HDL and non-HDL fractions. The isolated fractions were reconstituted to approximately the original serum volume and tested in culture. Scolex duplication was rarely observed in cultures supplemented with the non-HDL fraction. To assess whether tetrathyridia incorporate or recognize HDL, parasites were incubated with fluorescently labeled HDL. Strong fluorescence associated with the parasite surface was observed throughout the tetrathyridia (Fig. 3*C*).

### Knockdown of an HDL receptor homolog in *Mesocestoides vogae* suppressed asexual reproduction

We next sought to identify the parasite receptor responsible for recognizing HDL. In mammals, scavenger receptor class B type 1 (SR-B1), a member of the CD36 family, functions as a receptor for HDL. A putative CD36-family homolog was identified in *M. vogae* (XLOC_004160) (17). Previous transcriptomic analyses have reported that this gene is specifically expressed in the tetrathyridial stage (17).

During these experiments, we observed that scolex duplication frequency decreased with increasing time between parasite isolation from infected mice and serum supplementation (Fig. 4*A*). In parallel, expression of XLOC_004160 progressively declined following parasite isolation, accompanied by a reduction in HDL association with the parasite surface (Fig. 4*B, C*). These coordinated changes indicate a correlation between XLOC_004160 expression and serum responsiveness in tetrathyridia. To assess whether XLOC_004160 contributes to this phenotype directly, we performed RNAi-mediated knockdown. We found that dsRNA soaking enables RNA interference and successfully suppressed expression of the target gene in *M. vogae* tetrathyridia (Fig. S*1*). Upon serum supplementation, RNAi-mediated knockdown of XLOC_004160 significantly decreased scolex duplication frequency compared with controls treated with non-targeting dsRNA (Fig. 4*D*).

**Figure 4.**
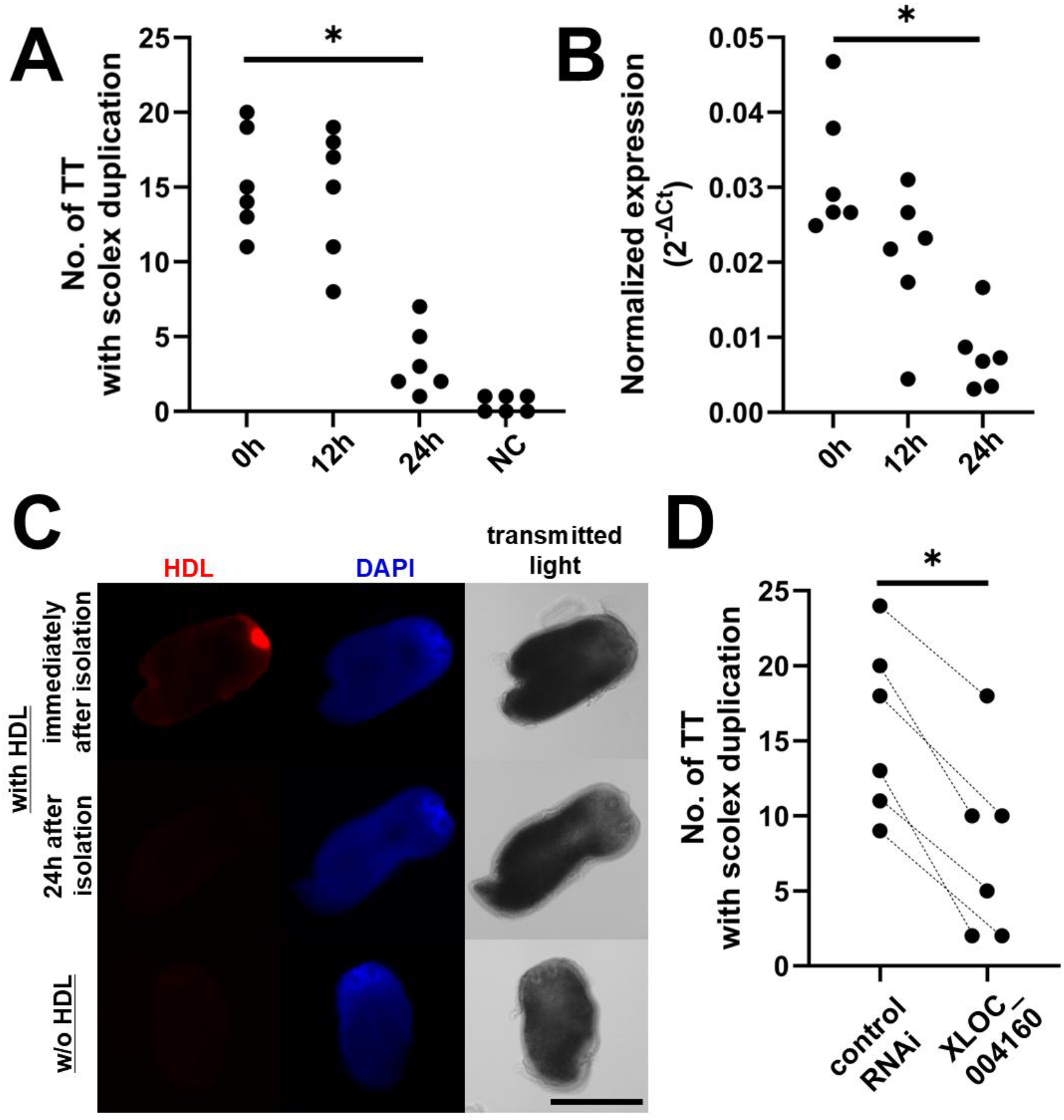
A CD36-family receptor homolog mediates HDL-dependent scolex duplication. (A) Frequency of scolex duplication as a function of time elapsed after tetrathyridia were isolated from the host prior to serum supplementation. (B) Expression of the CD36-family homolog XLOC_004160 after parasite isolation. (C) Association of fluorescently labeled HDL with tetrathyridia. (D) Effect of RNAi-mediated knockdown of XLOC_004160 on scolex duplication. Significance was evaluated using the two-sided Wilcoxon signed-rank test. \**P* < 0.05. Scale bar = 500 μm.

## Discussion

The present study demonstrates that host-derived HDL functions as an environmental cue that triggers the initiation of asexual reproduction in the larval stage of *M. vogae*. While mechanisms regulating reproductive activity after its onset have been relatively well studied (7–10), the host-derived signal that initiates the reproductive switch itself has remained unknown. By contrast, host-derived determinants that trigger the de novo onset of asexual reproduction, thereby initiating the transition from a nonreproductive to a reproductive state, have not been clearly identified in parasitic helminths.

In the peritoneal cavity, tetrathyridia are directly exposed to peritoneal fluid, and thus the use of the fluid for the present study would be preferable for reproducing physiological conditions. However, in the physiological state, the volume of peritoneal fluid that can be recovered from mice is extremely limited, making its collection technically challenging. Given that peritoneal fluid is largely derived from blood components that leak into the peritoneal cavity, serum was used in this study as a practical alternative. In a subset of infected mice from which sufficient peritoneal fluid could be obtained for measurement of HDL-cholesterol levels, we observed that the HDL-C concentration in the peritoneal cavity was approximately half of that in serum in all mice examined (Table S*1*). In this study, serum was diluted approximately twofold with culture medium, and scolex duplication was observed under the conditions (Fig. 1D). Thus, the HDL concentration under the experimental conditions, which should be close to that in peritoneal fluid, was sufficient to induce asexual reproduction in *M. vogae*.

Guinea pigs were reported to exhibit low serum HDL concentrations (15). In the present study, tetrathyridia survived for prolonged periods in the guinea pig peritoneal cavity but did not undergo asexual reproduction (Fig. 2*B*). HDL levels in guinea pig peritoneal fluid may be insufficient to reach the threshold required to initiate asexual reproduction. Consistent with this interpretation, *M. vogae* tetrathyridia were reported to undergo asexual reproduction in dogs, mice, jirds (*Meriones unguiculatus*) and cotton rats (*Sigmodon hispidus*) (11, 12, 18), all of which have relatively high serum HDL-C concentrations (216.9 ± 35.8 mg/dL, 97 ± 4 mg/dL, 72.64 ± 17.77 mg/dL, and about 80 mg/dL, respectively) (19–21). By contrast, rats have lower serum HDL-C concentrations (about 50 mg/dL) (19, 22), and asexual reproduction in rats was reported to be less pronounced than in mice (11). In addition, no scolex duplication was observed in guinea pigs (Fig. 2*B*), which have even lower HDL-C concentrations (approximately 6.0 ± 1.0 to 7.5 ± 1.5 mg/dL) than rats (19, 23). Furthermore, in the present study, supplementation with fetal bovine serum, which contains HDL-C at approximately one-thousandth the concentration found in mice (24), failed to initiate asexual reproduction (Fig. 1*D*). Collectively, these data suggest that HDL concentration may explain whether *M. vogae* is able to initiate asexual reproduction within a host and thus represents one factor underlying host suitability as an intermediate host. Asexual reproduction of *M. vogae* has also been reported in two additional host species, ground squirrels (*Citellus franklini*), and *Peromyscus maniculatus* (18). To the best of our knowledge, serum HDL-C concentrations have not been reported for either species. Determining their HDL-C levels would therefore help to clarify the relationship between HDL-C concentration and host suitability for *M. vogae*.

We identify XLOC_004160 as a putative HDL receptor that mediates the parasite response to host-derived HDL (Fig. 4*C*). However, direct evidence that the protein encoded by XLOC_004160 binds HDL is still lacking. We attempted forced expression of XLOC_004160 in mammalian cells. However, although the protein was expressed, it was not trafficked to the plasma membrane, preventing us from assessing whether its presence affects HDL binding at the cell surface. This is likely due to differences between cestodes and mammals in protein trafficking mechanisms, including membrane-targeting signals. At present, no cestode-derived cell lines amenable to genetic manipulation are available, making further investigation challenging. Detailed functional analysis of the cestode cell surface receptor will require the establishment of cestode cell lines and the development of genetic tools for these cells.

XLOC_004160 has been annotated as a sequence homolog of CD36 (17), which is a widely conserved member of the class B scavenger receptor family, present across a broad range of animals (25–27). *M. vogae* appears to have acquired the ability to undergo asexual reproduction through utilization of an existing gene rather than the evolution of a new gene. Not all cestode species reproduce asexually, but several species besides *M. vogae*, including *Taenia crassiceps* and *Sparganum proliferum*, are capable of asexual reproduction. These species are not monophyletic (28) and are distributed across multiple lineages within the cestode phylogeny, suggesting that the ability for asexual reproduction may have evolved several times independently. Notably, these other asexually reproducing species also possess XLOC_004160 homologs (Table *S2*). Whether they commonly utilize CD36 homologs in the initiation of asexual reproduction, representing parallel evolution via similar genetic changes, or whether each species employs distinct molecular systems to achieve asexual reproduction, representing convergent evolution through different molecular pathways, remains a question for future studies.

Genetic tools remain poorly developed in cestodes, and no studies have reported gene functional analysis using systemic knockout or knock-in approaches across cestodes (29, 30). At present, RNAi-mediated suppression of mRNA expression is virtually the only available approach for organismal-level phenotypic analysis in cestodes, but it has been reported in only a limited number of species, including *Moniezia expansa* and *Echinococcus multilocularis* (29–31). In the present study, we confirmed that RNAi-mediated gene silencing is also feasible in *M. vogae*, in addition to these cestode species. Because *M. vogae* can be readily maintained through asexual proliferation in the mouse peritoneal cavity, this cestode provides a tractable experimental model compared with other species that require multiple host animals. Combined with the RNAi technique reported here, this experimental system may provide a tractable platform to facilitate molecular and functional studies in cestodes.

Here, we provide evidence, not only in *M. vogae* but more broadly in parasitic helminths, of how parasites sense specific host-derived factors through defined parasite molecules to regulate transitions between life-cycle stages. Although *M. vogae* undergoes little asexual reproduction in the intestine (12), it proliferates extensively upon reaching the peritoneal cavity (11, 32). This observation can be interpreted as the parasite sensing differences in HDL concentrations to recognize that it has arrived at the appropriate host compartment for division. Our results demonstrate that cestode division is not a passive response to general nutrient availability, but is instead actively triggered by recognition of specific substances present in defined host compartments. Through such mechanisms, parasitic helminths can advance to the next life-cycle stage at the appropriate time and location within the host, ensuring coordinated development in response to the host environment.

## Materials and Methods

### Ethics statement

The present study was conducted in accordance with the recommendations in the Guide for the Care and Use of Laboratory Animals of Gifu University. The protocols were approved by the Committee on the Ethics of Animal Experiments of Gifu University (AG-P-N-20250167). Parasites were collected from mice after euthanasia by cervical dislocation. For all animal species, blood was collected by cardiac puncture under isoflurane inhalation anesthesia. Oral administration of parasites, cardiac puncture, and euthanasia by cervical dislocation were performed only by skilled researchers.

### Animals and parasites

Female BALB/c mice aged 8–12 weeks (Jackson Laboratory Japan, Yokohama, Japan) were used to passage *M. voga*e tetrathyridia. Serum samples were collected from female BALB/c mice aged 8–12 weeks (Jackson Laboratory Japan, Yokohama, Japan) or Slc:ICR mice aged 8–12 weeks (Japan SLC, Shizuoka, Japan). Female Slc:Hartley guinea pigs aged 6–8 weeks (Japan SLC) were used for experimental infection and serum collection. Blood samples were allowed to clot for 1 h and then centrifuged to separate serum. *Mesocestoides vogae* used in this study was originally isolated as described previously (32) and has since been maintained by serial passage in mice. Tetrathyridia were maintained in BALB/c mice via oral infection and harvested from their peritoneal cavities at least 120 days after infection. The harvested tetrathyridia were washed with phosphate-buffered saline (PBS) five times.

### Evaluation of serum-induced scolex duplication in tetrathyridia

The basic medium consisted of RPMI-1640 medium (Sigma-Aldrich, St. Louis, MO, USA) containing 20% heat-inactivated fetal bovine serum (56 °C, 30 min) and 1× Antibiotic-Antimycotic Mixed Stock Solution (Nacalai Tesque, Kyoto, Japan). All experiments were conducted in 12-well plates with 25 tetrathyridia bearing a single scolex per well. To each well containing 400 µL of basic medium, an additional 500 µL of non-heat-inactivated BALB/c mouse, guinea pig, or fetal bovine serum was added, resulting in a total volume of 900 µL per well, followed by incubation at 37 °C for 7 days in a 5% CO_2_ incubator. After incubation, the number of tetrathyridia exhibiting scolex duplication was counted using a stereomicroscope.

### Evaluation of scolex duplication induced by ultrafiltration fractions of serum

Serum ultrafiltration was performed using 500 µL of BALB/c mouse serum and an Ultra Centrifugal Filter, 100 kDa MWCO (APRO Science, Tokushima, Japan) at 14,000 × *g* for 30 min at room temperature. The retained and flow-through fractions were each adjusted to the original serum volume with basic medium for subsequent tetrathyridium culture experiments. Culture assays were performed in 12-well plates containing 25 single-scolex tetrathyridia per well. Each well received 400 µL of basic medium with 500 µL of adjusted, retained, flow-through fraction, or BALB/c mouse serum. After incubation for 7 days at 37 °C in 5% CO_2_, the frequency of scolex duplication was determined under a stereomicroscope.

### Evaluation of scolex duplication induced by high-density lipoprotein

BALB/c mouse serum-derived HDL was purified using HDL Purification Kit (Ultracentrifugation Free) (Cell Biolabs, San Diego, CA, USA) according to the manufacturer’s instructions. Purified HDL was recovered in the extraction solvent supplied with the kit. To remove this solvent, an additional ultrafiltration step was performed using an Ultra Centrifugal Filter, 100 kDa MWCO (APRO Science) at 14,000 × *g* for 30–40 min at room temperature, until the retentate was reduced to a minimal volume. HDL purified from 1 mL of serum was retained on the filter and then resuspended in 400 µL of basic medium, and the resulting preparation was directly used for tetrathyridium culture. Culture conditions and evaluation methods for scolex duplication were as described above.

HDL, VLDL, LDL, and CMs were separated from BALB/c or Slc:ICR mouse serum according to the method of Hatch et al. (33). Because the isolated lipoprotein fractions were present in the density gradient medium used for ultracentrifugation, the solvent volume was reduced to a minimal volume by ultrafiltration, as described above. Among lipoproteins separated from 4 mL of serum, HDL was then resuspended separately, whereas the other lipoprotein fractions were pooled and resuspended together in 400 µL of basic medium. The resulting preparations were directly used for tetrathyridium culture. Culture conditions and evaluation methods for scolex duplication were as described above.

### Binding of fluorescently labeled HDL to tetrathyridia

Lipoprotein-containing components in Slc:ICR mouse serum were first labeled with the lipophilic fluorescent dye 1,1′-dioctadecyl-3,3,3′,3′-tetramethylindocarbocyanine perchlorate (DiI) according to the method described by Pitas et al. (34). Subsequently, HDL was purified from the labeled serum using the procedure described above. Tetrathyridia were cultured in medium containing fluorescently labeled HDL for 12 h, fixed with 4% paraformaldehyde (PFA), permeabilized with 0.1% Triton X-100, and stained with DAPI. Samples were observed using an all-in-one fluorescence microscope (Keyence, Osaka, Japan).

### RNA interference (RNAi)

Total RNA was extracted from 50 tetrathyridia using TRIzol Reagent (Thermo Fisher Scientific, Waltham, MA, USA). First-strand cDNA was synthesized from the total RNA using an oligo(dT) primer and the PrimeScript 1st Strand cDNA Synthesis Kit (Takara Bio, Shiga, Japan), according to the manufacturer’s instructions. Using this cDNA as a template, the XLOC_004160 fragment was amplified by polymerase chain reaction (PCR) with Tks Gflex polymerase and primers XLOC_004160 F and R (primer sequences and PCR conditions are listed in Table S1; the same applies hereafter). The PCR product was ligated into the pTA2 vector using the TArget Clone - Plus-kit (Toyobo, Osaka, Japan) and transformed into DH5α competent cells (Takara Bio).

Plasmids were isolated from the resulting colonies, and the inserted sequence was confirmed by sequencing to be identical to the sequence registered in WormBase Parasite (https://parasite.wormbase.org/Mesocestoides_corti_prjeb510/Gene/Summary?g=XLOC_004160;r=MCOS_contig0001049:9194-11936).

To generate templates for single-stranded RNA (ssRNA) synthesis, a partial region of the target sequence was amplified by PCR using the plasmid as a template and Tks Gflex DNA polymerase (Takara Bio), with two sets of primers (XLOC_004160 ssRNA F/R T7 and F T7/R; Table S*1*). PCR products were purified using the NucleoSpin Gel and PCR Clean-up kit (Macherey-Nagel, Düren, Germany) according to the manufacturer’s instructions.

To allow annealing and generate dsRNA, the two complementary ssRNAs were first incubated at 65 °C for 5 min, mixed in equal amounts (5.0 × 10^4^ ng each), and then incubated at 70 °C for 10 min. The temperature was subsequently decreased by 0.5 °C every 13 s until reaching 25 °C.

As a negative control, dsRNA targeting a tomato (*Solanum lycopersicum*) chloroplast-specific ribosomal protein cDNA (GenBank accession No. AY568722) was generated from RNA extracted from commercially purchased tomato fruit using the same procedure as described above. PCR conditions and primer sequences are provided in Table S*1*.

We added 1.0 × 10^5^ ng of dsRNA targeting XLOC_004160 or an unrelated tomato gene (negative control) to 400 µL of basic medium, and 40 tetrathyridia with a single scolex were incubated for 12 h. After incubation, 25 tetrathyridia were collected without any special selection and transferred to 400 µL of fresh basic medium supplemented with 500 µL of BALB/c mouse serum. The tetrathyridia were cultured for 7 days, after which the frequency of scolex duplication was determined. The remaining 15 tetrathyridia were subjected to total RNA extraction as described above and used for subsequent experiments.

To confirm the effect of RNAi, total RNA extracted from the remaining 15 tetrathyridia was used to assess the mRNA level of XLOC_004160 by real-time PCR under the conditions shown in Table S*1*. Real-time PCR reactions were carried out in a Thermal Cycler Dice Real Time System III (Takara Bio) using TB Green Premix Ex Taq (Tli RNaseH Plus) (Takara Bio). Two *M. vogae* transcripts, high molecular weight tropomyosin 1 (HMW-TPM1) and glyceraldehyde-3-phosphate dehydrogenase (GAPDH), were used as endogenous reference genes, as in a previous study (38). Differences in the obtained threshold cycle (Ct) values between samples were calculated using the ΔCt method.

### Basic local alignment search tool (BLAST)-based search for XLOC_004160 homologs

The predicted amino acid sequence of XLOC_004160 (Table S2) was used as a query for BLASTP searches against cestode sequences using the BLAST program available at the NCBI BLAST website (https://blast.ncbi.nlm.nih.gov/Blast.cgi). All hits with E-values less than 1.0 × 10^−10^ were collected. Putative functions of the proteins inferred from the retrieved amino acid sequences were evaluated using the PANTHER classification system (version 19.0).

### Statistical analysis

All statistical analyses were performed using R (version 4.4.0) (39). Comparisons between samples were conducted using a two-sided Wilcoxon signed-rank test, and *P* values were adjusted for multiple comparisons using the Holm method.

## Declaration of Generative AI and AI-Assisted Technologies

ChatGPT (OpenAI, GPT-5.5) was used solely for grammar and language editing during the preparation of this manuscript.

## Author Contributions

T.S. and Y.T. designed research; T. S., K.H. and K.N. performed research; T.S. analyzed data; Y.T. provided funding and supervision; and T.S., K.H., K.N. and Y.T. wrote the paper.

## Acknowledgments

This research was supported by the Japan Society for the Promotion of Science, Grant-in-Aid for Scientific Research (A) 23H0447 and for Early-Career Scientists 25K18361. We thank Robin James Storer, PhD, from Edanz (https://jp.edanz.com/ac) for editing a draft of this manuscript.

## Data availability

Study data are included in the article and supporting information.

